# A highly contiguous genome for the Golden-fronted Woodpecker (*Melanerpes aurifrons*) via a hybrid Oxford Nanopore and short read assembly

**DOI:** 10.1101/2020.01.03.894444

**Authors:** Graham Wiley, Matthew J. Miller

## Abstract

**Background:** Woodpeckers are found in nearly every part of the world, absent only from Antarctica, Australasia, and Madagascar. Woodpeckers have been important for studies of biogeography, phylogeography, and macroecology. Woodpeckers hybrid zones are often studied to understand the dynamics of introgression between bird species. Notably, woodpeckers are gaining attention for their enriched levels of transposable elements (TEs) relative to most other birds. This enrichment of TEs may have substantial effects on woodpecker molecular evolution. The Golden-fronted Woodpecker (*Melanerpes aurifrons*) is a member of the largest radiation of New World woodpeckers. However, comparative studies of woodpecker genomes are hindered by the fact that no high-contiguity genome exists for any woodpecker species.

**Findings:** Using hybrid assembly methods that combine long-read Oxford Nanopore and short-read Illumina sequencing data, we generated a highly contiguous genome assembly for the Golden-fronted Woodpecker. The final assembly is 1.31 Gb and comprises 441 contigs plus a full mitochondrial genome. Half of the assembly is represented by 28 contigs (contig N50), each of these contigs is at least 16 Mb in size (contig L50). High recovery (92.6%) of bird-specific BUSCO genes suggests our assembly is both relatively complete and relatively accurate. Accuracy is also demonstrated by the recovery of a putatively error-free mitochondrial genome. Over a quarter (25.8%) of the genome consists of repetitive elements, with 287 Mb (21.9%) of those elements assignable to the CR1 superfamily of transposable elements, the highest proportion of CR1 repeats reported for any bird genome to date.

**Conclusion:** Our assembly provides a useful tool for comparative studies of molecular evolution and genomics in woodpeckers and allies, a group emerging as important for studies on the role that TEs may play in avian evolution. Additionally, the sequencing and bioinformatic resources used to generate this assembly were relatively low-cost and should provide a direction for the development of high-quality genomes for future studies of animal biodiversity.

## Introduction

Because of their near-global distribution, woodpeckers are often used as model systems in biogeographic and phylogeographic studies [1–5], as well as macroecological studies [6–9]. In North America, four woodpecker hybrid zones have also been studied for insights into avian speciation. These include flickers [10–15], sapsuckers (*Sphyrapicus;* [10–15], Nuttall’s/Ladder-back woodpeckers [16] and Red-bellied/Golden-fronted woodpeckers [10–15]. More recently, woodpeckers have gained attention for the high amount of repetitive DNA found in their genomes relative to other bird taxa [17–19], the result of high levels of genome-wide transposable elements (TEs), which are scarce in most bird genomes [20]. Manthey et al. [21] surveyed several woodpecker genomes, and found that TEs make up 17–31% of woodpecker genomes, compared to <10% for other bird species. Increasingly, researchers have suggested that TEs may play a critical role in driving avian evolution [20,22]. However, only one woodpecker genome (*Picoides* [*Dryocopus*] *pubescens*) has been published, limiting the ability to more fully understand the architecture of TE evolution in woodpeckers, and more generally for comparative genomics in the group.

The genus *Melanperpes* represents the largest radiation of New World woodpeckers (Aves: Picidae) [4]. *Melanerpes* woodpeckers are found almost everywhere where forest occurs in the Americas. Various species range continuously from southern Canada to Argentina, with three species occurring in the West Indies. Members of the Red-bellied/Golden-fronted Woodpecker species complex (*Melanerpes aurifrons/carolinus*) are notable for the discord between plumage variation and phylogenetic structure, especially in Mexico and northern Mesoamerica, where various races show considerable plumage variation despite a lack of phylogeographic variation [4,23], see also [24].

Here, we provide a high-quality, highly contiguous, reference genome for the Golden-fronted Woodpecker (*Melanerpes aurifrons*; Figure 1), which we expect to be foundational for research in avian hybrid zone dynamics, the evolution of reproductive isolation, and the role of TEs and other repetitive elements in driving molecular evolution and speciation in woodpeckers and other birds. Additionally, we believe our workflow will be of general interest to researchers looking to develop high quality reference genomes for non-model birds and other vertebrates. Our genome required only ∼46X coverage of long-read Oxford Nanopore and ∼52X coverage of short-read Illumina sequence. Furthermore, after troubleshooting the various steps in our workflow, we were able to go from raw data to completed assembly in less than two weeks, using a compute cluster with 72 CPU cores, 384 GB of RAM, and two NVIDIA Tesla P100 GPU accelerators. These sequencing and computational requirements should be fairly accessible to most research groups with modest budgets.

**Figure 1.**
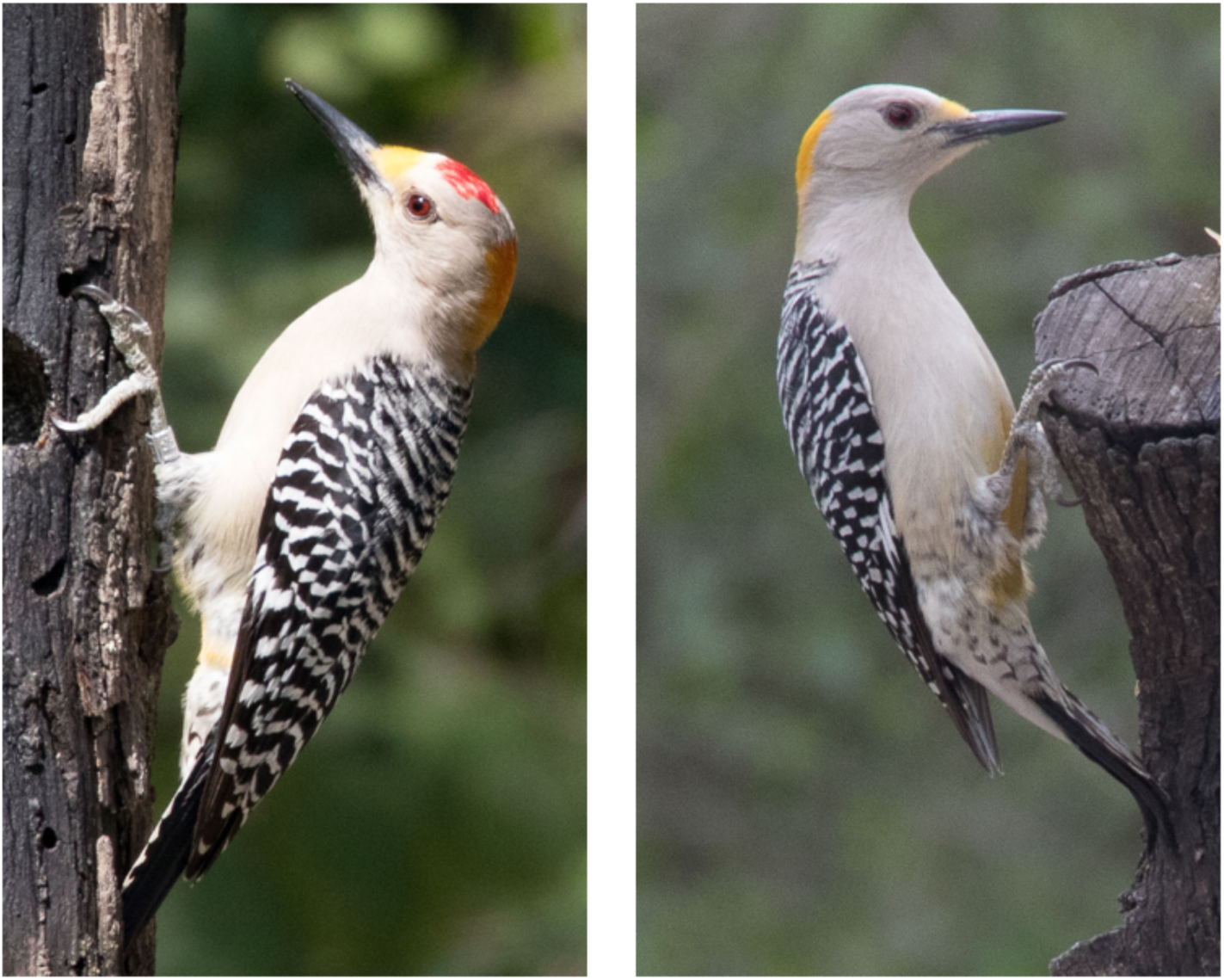
Male (left) and female (right) Golden-fronted Woodpecker (*Melanerpes aurifrons*). Photos by Bettina Arrigoni, cropped, and used under CC BY 2.0 license. Original photos available at: https://flickr.com/photos/69683857@N05/39849351035 and https://flickr.com/photos/69683857@N05/26752528708.

## Methods

### Specimen collection and DNA extraction

We collected an adult female Golden-fronted Woodpecker (*Melanerpes aurifrons*) in March 2019 in Dickens, Texas. This specimen and all associated genomic resources have been archived as a museum voucher at the Sam Noble Oklahoma Museum of Natural History (specimen number 24340, tissue number SMB 682). This specimen is registered with NCBI as Biosample SAMN13719207. Scientific collecting was done under the following permits: Texas Scientific Collecting Permit SPR-0916-229, US Migratory Bird Treaty Act permit MB02276C-0, and University of Oklahoma IACUC permit R15-016A. Approximately ten milliliters of whole blood were transferred immediately to blood tubes coated with EDTA as an anticoagulant and kept cool on wet ice for 6 hours. Upon return to the lab, blood was aliquoted to several tubes and stored at −20°C. We extracted high molecular weight DNA from a single aliquot using the Nanobind CBB Big DNA Kit (Circulomics, Inc). We followed the manufacturer’s high molecular weight (50–300+ kb) DNA protocol for cultured mammalian cells, with the exception that we diluted 10 uL of whole woodpecker blood into 190 uL of fresh PBS as sample input. This was done per recommendation of the manufacturer (personal communication) given that avian red blood cells are nucleated. We performed three rounds of HMW DNA extraction, with each extraction being used as input for an Oxford Nanopore sequencing library. For the initial MinION nanopore sequencing run, we treated the HMW DNA with Short Read Eliminator Kit (Circulomics, Inc.), following the manufacturer’s instructions. For the PromethION sequencing runs, this kit was not used in order to maximize DNA concentration for downstream library construction. For Illumina sequencing, we performed a standard genomic DNA extraction using the Quick-DNA miniprep kit (Zymo Research). Again, 10 uL of blood was diluted into 190 uL of PBS before following the manufacturer’s protocol.

### Whole genome library preparation and sequencing

We first generated Oxford Nanopore long reads in house on the MinION device using the SQK-LSK109 library preparation kit followed by sequencing on a R9.4.1 flow cell per the manufacturer’s instructions (Oxford Nanopore Technologies). After observing low yield, we sent our remaining two HMW extracts to the UC Davis Core Lab to run on two respective PromethION flow cells. For the first PromethION run a single library was prepared, loaded, and run for 48 hours. After limited yield in that run, for the second PromethION flow cell, the HMW DNA was sheared using a Megaruptor (Diagenode, Inc.) set to a 50 kb target prior to library preparation. Two libraries were made such that the flow cell ran for 24 hours, was flushed with a nuclease wash, and a second library was loaded for the final 24 hours of run time. To generate the Illumina sequencing library, we used the NEBNext Ultra II FS DNA library kit (New England Biolabs, Inc); the initial enzymatic shearing step was accomplished via 10 minutes of incubation at 37°C, after which we followed the manufacturer’s instructions. The library was indexed using NEBNext Multiplex Oligos for Illumina unique dual index kit (New England Biolabs, Inc). This library was run on one lane of a NovaSeq S-Prime flow cell at the Oklahoma Medical Research Foundation Clinical Genomics Center.

### Read QC and trimming

Raw Nanopore reads (fast5) were converted into fastq format using Oxford Nanopore’s proprietary base-calling software guppy v.3.4 (https://community.nanoporetech.com). We evaluated Nanopore read quality in NanoPlot, and filtered reads (Q>7, length>10,000) using NanoFilt; both are part of the NanoPack distribution [25]. We filtered and trimmed Illumina reads using the standard settings in the bbduk program, which is part of BBTools v38.00 (Bushnell 2019).

### Genome size estimation

We used jellyfish v2.2.3 [26] to count the frequency of three distinct *k*-mers (17-mers, 26-mers, and 31-mers) in our trimmed Illumina sequencing reads. We plotted the resulting histogram file to establish the peak coverage at each *k*-mer. We then used the following formula from Liu et al [27]to estimate genome size:

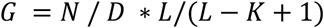

where *G* equals genome size, *N* equals the total number of base pairs in all reads, *D* equals the expected *k*-mer coverage depth, *L* equals the average read length, and K equals the *k*-mer size.

### Genome assembly

We assembled an initial draft assembly using flye v2.6 [28,29].

We corrected errors in that assembly using three iterations of racon v1.4.7 [30], which uses the original nanopore long reads to correct the assembly consensus output. Subsequently, we polished the genome with Illumina reads using two iterations of polishing using pilon v1.23 [31]; we used minimap2 v2.17 [32] to map the Illumina short reads to the draft genome to generate the assembly for input for pilon. Finally, we removed redundant haploid and low coverage contigs from the polished assembly using purge_haplotigs v1.1 [33]. That workflow begins by generating a read depth histogram to establish coverage cutoff levels for low coverage, high coverage, and the midpoint in coverage between haploid and diploid coverage. Subsequent steps remove contigs if they fall outside this range. To prevent interference from repeat elements, a bed file containing repetitive element coordinates was provided to purge_haplotigs (see below). At each step, we evaluated the completeness of the genome assembly via BUSCO v3 benchmarking [34] using the aves_odb9 dataset, which contains 4915 near-universal single-copy orthologs.

### Identifying repetitive elements and annotating the mitochondrial genome

We estimated the portion of our genome comprised of repetitive elements using RepeatMasker v4.0.9p2. [35]. To ensure avian-specific annotations we utilized the ‘chicken’ species entry in the RepeatMasker database. The output of this was converted to a bed file for use in purge_haplotigs (see above).

To identify a contig that may represent the mitochondrial genome, we performed BLASTn against an existing sequence of *M. aurifrons* NADH-dehydrogenase subunit II (GENBANK: MF766655) against our draft assembly, which recovered a single contig. Using that contig as a draft mitochondrial genome, we assembled all mitochondrial genes using the MitoAnnotator web server [http://mitofish.aori.u-tokyo.ac.jp/; 36]. As an additional metric to evaluate the robustness of our assembly, we assessed the quality of that assembly by comparing all protein coding gene sequences to an annotated mitochondrial genome of the Great Spotted Woodpecker (*Dendrocopos major*, GENBANK: NC028174) and we confirmed proper secondary structure for all tRNAs using the tRNAscan-SE 2.0 web server [http://lowelab.ucsc.edu/tRNAscan-SE/index.html; 37]. Finally, we did a Google Scholar search to look for other avian genomes published in 2019 to compare our results to other high-quality wild bird reference genomes.

### Reference assisted pseudomolecule scaffolding

We generated pseudomolecule chromosomal scaffolds using a reference genome assisted approach in RaGOO v1.1 [38]. RaGOO attempts to cluster, order, and orient assembly contigs based on a Minimap2 [32] alignment of those contigs to a reference genome. There is no chromosomal-level genome available for woodpeckers and allies (Order: Piciformes) and all bird species with chromosomal-level genomes are roughly similar in phylogenetic distance [39]. Therefore, we chose to use the Anna’s Hummingbird chromosome-assembled genome [Calypte anna bCalAnn1_v1.p; GCA_003957555.2; 40] as our reference. To evaluate the validity of our reference-guided scaffold we measured genome synteny by aligning the RaGOO-scaffolded assembly to the Anna’s Hummingbird reference genome as well as to two additional chromosomal bird genomes: Zebra Finch (Taeniopygia guttata, GCA_008822105.1) and Kakapo (*Strigops habroptila*, GCA_004027225.1). We used the nucmer module in MUMMER v4.0.0b2 [41] to perform the alignments which were subsequently filtered using MUMMER’s delta_filter module with many-to-many alignments allowed, minimum alignment length equal to 500, and minimum alignment identity equal to 80%. We filtered this output to only select mapped clusters equaling at least 3000 base pairs in the RaGOO (query) assembly. To visualize genome synteny we generated circos plots in Circa (http://omgenomics.com/circa); to improve visual clarity, only chromosomes 1 through 14 (including 4A, 4B and 5A) and the Z and W chromosomes were used.

## Results and Discussion

### Sequencing run statistics and genome assembly results

We generated a total of 3.9 M Oxford Nanopore reads for a total of 63.99 Gb of long read sequence data (Table 1). The initial MinION sequencing run produced only limited data but had longer reads than either the unsheared or sheared HMW runs on the PromethION (Table 1). Megaruptor shearing resulted in slightly lower median read length but increased read length N50. Comparing our two PromethION runs, HMW shearing, along with reloading a fresh library after the initial 24-hour runtime, increased the number of reads recovered by 18% and total base pairs yielded by 40%. These results, along with the slightly higher read length N50, suggest that users should consider HMW DNA shearing prior to Oxford Nanopore library preparation for metazoan whole genome assemblies.

**Table 1.** Run statistics for three Oxford Nanopore sequencing runs.

| Sequencing Run | Reads | Median Read Length | Read Length N50 | Median Read Qual | Total Bases |
| --- | --- | --- | --- | --- | --- |
| MinION Run | 101,989 | 15,105 | 39,203 | 10.5 | 2.29 X 10<sup>9</sup> |
| PromethION Run 1 | 1.77 X 10<sup>6</sup> | 9,478 | 30,742 | 10.6 | 27.43 X 10<sup>9</sup> |
| PromethION Run 2 | 2.09 X 10<sup>6</sup> | 9,178 | 34,314 | 10.5 | 34.27 X 10<sup>9</sup> |

Our initial flye assembly resulted in 1519 contigs which were organized into 1456 scaffolds with 41X mean coverage. The total length of the assembly was 1.35 Gb with a contig L50 of 14.5 Mb and a scaffold L50 of 15.9 Mb. The largest contig was 4.28 Mb. Fifty percent of the assembly was recovered in just 33 contigs (L50) – those contigs were at least 14.5 Mb in size (550). Ninety percent of the assembly was recovered in 136 contigs (L90) all of which were at least 1.4 Mb in size (N90). Our final genome assembly resulted in 441 ungapped contigs with an estimated genome size of 1.309 Gb, plus a complete circularized mitochondrial genome (N = 16,844). The largest contig was 46.8 Mb. Contig L50 was 28 contigs and contig N50 was 16 Mb.

Our Illumina sequencing generated a total of 490.7 M paired end reads for a total of 74.1 Gb of short read sequence data. After bbduk trimming, we retained 71.2 Gb of Illumina data from 487.8 M reads. Histograms for k-mer coverage (Supplementary Figure S1) suggest an average k-mer depth ranging from 46 (*k*-mer = 17 bp) to 41 (*k*-mer = 31 bp). This results in an estimated genome size of 1.38 – 1.40 Gb, depending on the *k*-mer value (Table 2). For resulting analyses, we consider the genome size estimate from the 31-mer analysis (1.38 Gb) to be the best estimate for our genome. This suggests that our assembly recovered 94.2% of the true Golden-fronted Woodpecker genome. A recent review suggests that on-average, bird genomes generated with short reads are 11–29% incomplete [42]. Given our genome size estimate, our Oxford Nanopore reads were at an average depth of 43.4, while our trimmed Illumina reads were at an average sequencing depth of 51.6.

**Table 2.** Results of genome size estimate using *k*-mer analysis.

| *k*-mer size | *k*-mer coverage | Estimated genome size |
| --- | --- | --- |
| 17 | 46 | 1.377 Gb |
| 26 | 42 | 1.404 Gb |
| 31 | 41 | 1.379 Gb |

### Racon correction eliminates gaps and merges small contigs

Consensus correction using racon eliminated gaps in scaffolds, meaning that all fragments were ungapped contigs. Racon greatly reduced the number of contigs, but this was apparently mostly achieved by merging small contigs with overlapping regions from larger ones. Thus, racon had the effect of slightly reducing assembly size with only slight increases in contig L50, L90, N50, and N90 (Table 3). The largest effect of racon correction was observed in the first iteration, by the third round, little change was observed.

**Table 3.**
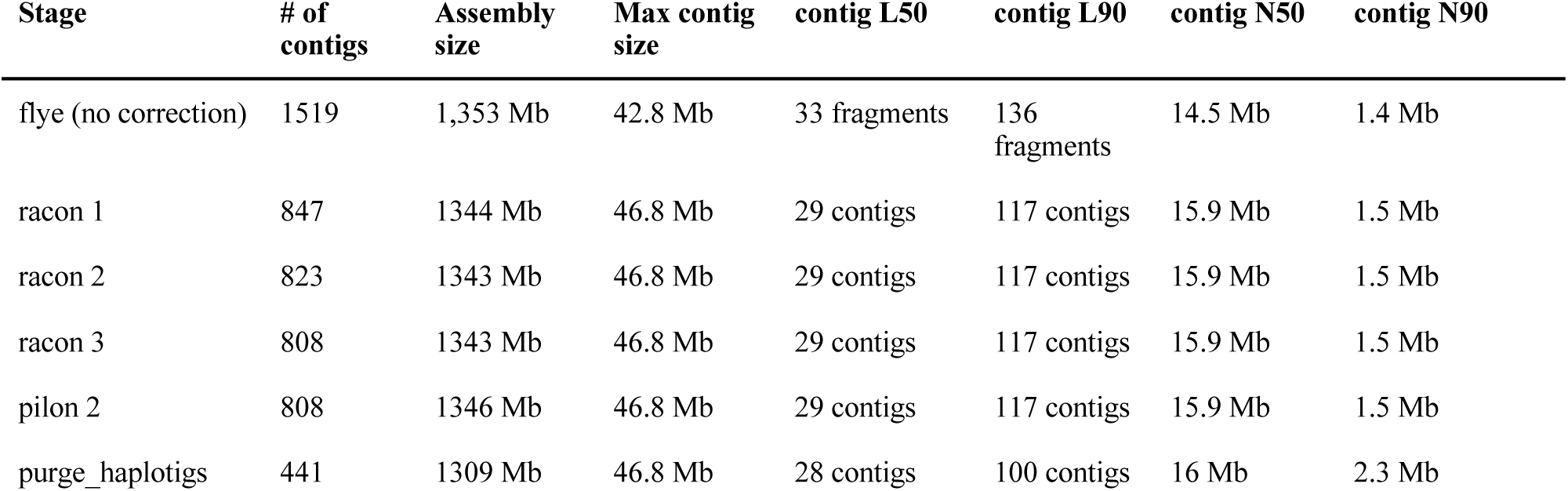
Assembly contiguity statistics for the initial assembly and after multiple iterations of consensus correction.

### Polishing and haplotig purging improve genome accuracy

Whereas racon improved the contiguity of the assembly, it achieved only modest improvement in BUSCO benchmarking (Table 4). On the other hand, while pilon had essentially no effect on assembly contiguity (Table 3), it greatly improved BUSCO benchmarking, which we interpret as an estimation of correctness of genome-wide base calls. A second iteration of pilon resulted in only negligible improvement in BUSCO benchmarking. Haplotig purging had negligible impact on the overall completeness calculations, with a 0.2% reduction in duplications and a 0.1% increase in missing orthologs. Instead, purge_haplotigs identified 366 contigs as redundant, resulting in a more accurate genome representation while maintaining the overall genome assembly. Our final assembly recovered as complete 92.6% of the BUSCO orthologs.

**Table 4.** BUSCO summarized benchmarking at various stages of genome assembly.

| Stage | Complete (Single, Duplicate) | Fragmented | Missing |
| --- | --- | --- | --- |
| flye (no correction) | 76.2% (74.7%, 1.5%) | 10.6% | 13.2% |
| racon 3 | 76.5% (75.1%, 1.4%) | 11.0% | 12.5% |
| pilon 1 | 92.6% (90.8%, 1.8%) | 4.5% | 2.9% |
| pilon 2 | 92.7% (90.8%, 1.9%) | 4.5% | 2.8% |
| purge_haplotigs | 92.6% (90.9%, 1.7%) | 4.5% | 2.9% |

### Repetitive elements in the assembly

RepeatMasker estimates that 305 Mb (25.8%) of our assembly is comprised of repetitive elements. Nearly all of these elements (93.9%) – a total of 287 Mb (21.9% of the genome) – are part of the CR1 transposable element (TE) superfamily.

Birds have smaller genomes than most other tetrapods [43], likely due to the metabolic cost of flight [44]. The streamlining of avian genomes appears to be driven by a substantial reduction in the frequency of TEs in the genome [20]. For most tetrapods, TEs represent a sizeable portion of the genome. For example, in mammals TEs typically account for 1/3 to 1/2 of the entire genome [45], while in birds’ closest living relatives, the crocodiles, TEs represent about 1/3 of the genome [46]. In contrast, birds have substantially lower frequency of TEs – typically ranging from 4–10% of the total genome size [19]. Woodpeckers and allies are the exception. Manthey et al. [21] demonstrated that TE occurred at higher frequency across several genera of woodpeckers (17–30%), due principally to expansion of CR1 TEs (15–17% of total TEs). Our finding of nearly 22% attributable to CR1 TEs represents a higher proportion of a woodpecker genome than any reported by Manthey et al. Because Manthey et al.’s technical approach likely under-estimates TE content, direct comparisons should be considered tentative, but our result does represent the highest CR1 proportion reported for any avian genome.

### Mitogenome assembly and annotation results

The mitochondrial genome assembly (Figure 2) recovered all 13 protein-coding genes, 12S and 16S ribosomal RNAs, and 22 tRNAs, all in the standard order observed in other woodpeckers and allies [47,48]. Pairwise alignment of each of these features shows all features starting and ending coincident with homologous features in the Great Spotted Woodpecker mitochondrial genome, except in a couple of cases where our initial annotation failed to properly assess protein-coding genes with incomplete (-T) stop codons, which are common in vertebrate mitochondrial genomes [49]. In nearly all cases where this occurred, the adjacent 3’ base would result in a valid stop codon, so this “error” seems to be a limitation of the MitoAnnotator algorithm rather than a measure of error in our mitogenome assembly. Overall, the quality of our mitogenome assembly provides independent evidence to suggest that racon consensus correction plus pilon short read polishing is able to correct most errors in long read genome assemblies.

**Figure 2:**
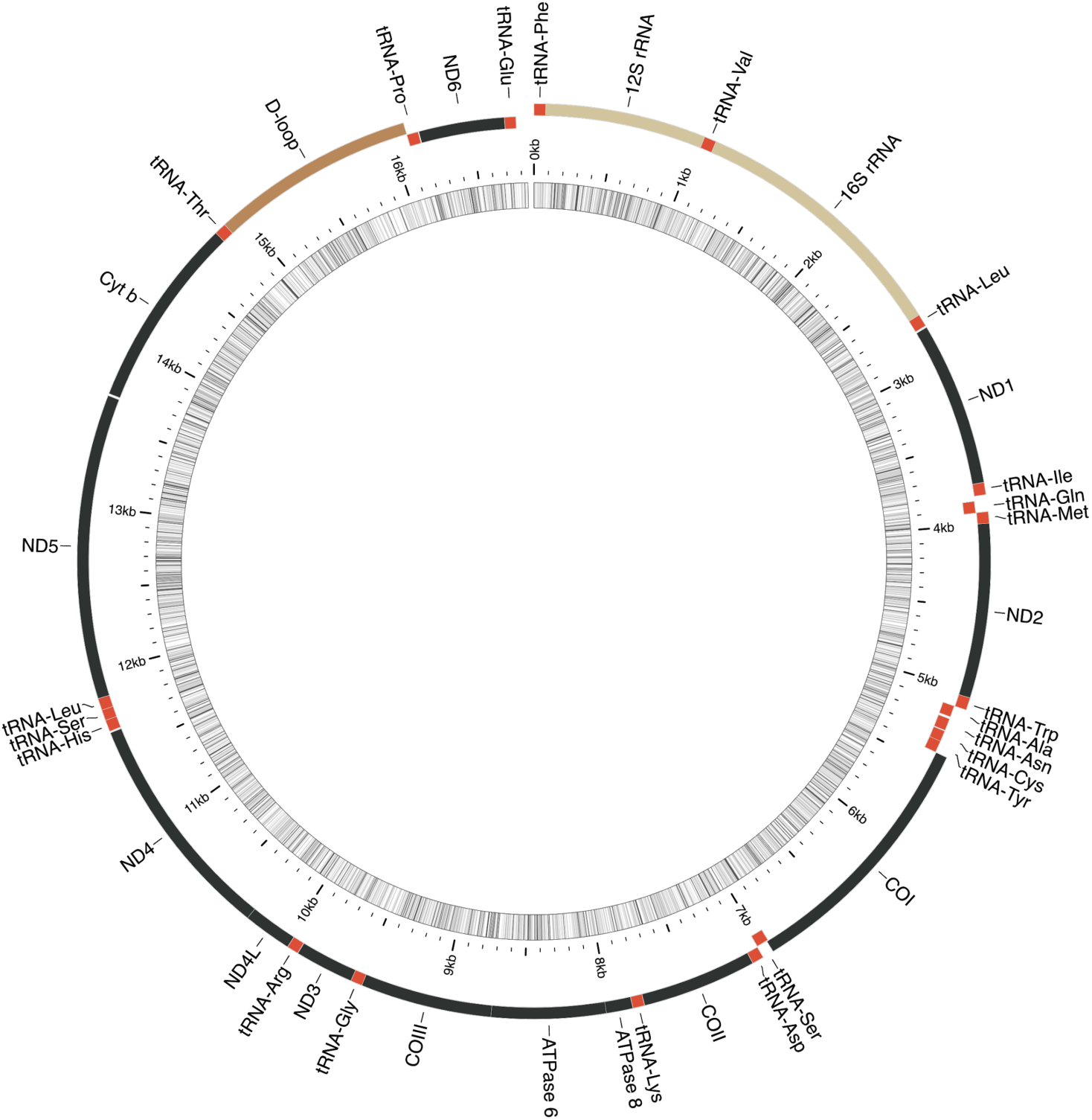
Circularized annotated mitochondrial genome assembly for the Golden-fronted Woodpecker (*Melanerpes aurifrons*). Figure generated by MitoAnnotator [36].

### Synteny with other avian genomes and the usefulness of reference-based scaffolding

In general, synteny plots show that most of our Anna-s Hummingbird guided reference-based scaffolded pseudo-chromosomes had largely one-to-one synteny plots with chromosomes of all three bird genomes evaluated (Figure 3). In some instances, the chromosomal numbers do not agree. For example, our pseudo-chromosome 5A maps almost exclusively to the Kakapo chromosome 4. This is likely to be simply the result of differences between chromosome numbering conventions. Both the Zebra Finch and Anna’s Hummingbird synteny plot show that a portion of those birds’ W chromosomes map to our Golden-fronted Woodpecker pseudo-chromosome 3, suggesting that this pseudo-molecule may be misassembled. We agree with Alonge et al. [38] that reference-guided assemblies will be generally useful for comparative genomics studies of animal biodiversity, such as to map genome-wide phylogenetic markers, or to map the locations of structural variants and/or *F*_*st*_ outliers, etc. However, given the relatively low number of contigs in our assembly, researchers may prefer to do those analyses on the 441 unscaffolded contigs rather than on the pseudo-chromosomes.

**Figure 3:**
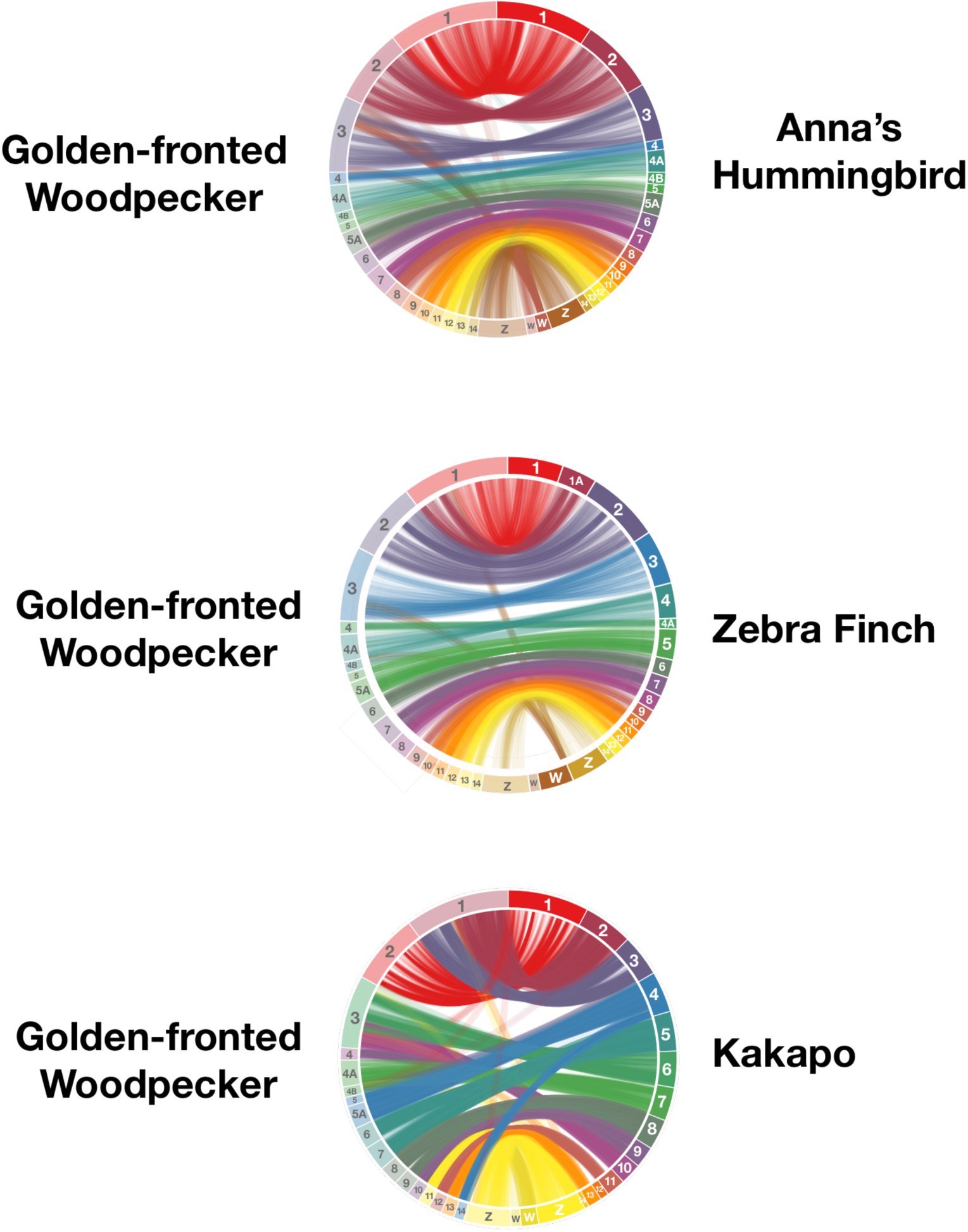
Ideograms showing synteny between Golden-fronted Woodpecker pseudo-chromosomes and chromosomes from three different bird species.

### Comparisons to recently published avian genomes

There is increasing interest in obtaining reference genomes for wild bird research. While a full chromosome level assembly would be ideal, many research questions would benefit from a highly contiguous but less-than-chromosome assembly. How should research groups with limited resources go about generating such genomes? Comparing assembly statistics for some recently published avian genomes (Table 5), we find that hybrid (PacBio or Oxford Nanopore long reads + Illumina short reads) assemblies are generally more contiguous than assemblies generated with only Illumina sequencing, including those with third generation Illumina libraries (e.g. 10X and Hi-C). And our genome may be more complete than many other published genomes [42]. As de novo genome assembly algorithms such as flye [29], RedBean [50], Shasta (https://chanzuckerberg.github.io/shasta/), and Canu [51] that are capable with dealing with noisy Oxford Nanopore data continue to be developed, we expect to see more researchers pursuing in-house genome assemblies for their own studies of animal biodiversity genomics.

**Table 5:** Comparison of various genome assembly statistics for recently published wild bird genomes. * N50 scaffold not reported for assemblies comprised only of ungapped contigs. ** Statistics only reported for scaffolded contigs.

| Species | Reference | Method | Assembly size | # of contigs/# of scaffolds | N50 contig | N50 scaffold | BUSCO complete | % Repeats |
| --- | --- | --- | --- | --- | --- | --- | --- | --- |
| *Pavo cristatus* (Galliformes) | [52] | Nanopore (low coverage) + short read | 932 Mb | 685,241/ 15,025 | 14.7 kb | 0.23 Mb | Not reported | 7.3% |
| *Melospiza melodia* (Passeriformes) | [53] | Paired end + HiC reads | 978 Mb | Not reported | 31.7 kb | 5.6 Mb | 87.5% | 7.4% |
| *Rhegmatorhina melanosticta* (Passeriformes) | [54] | Paired end + 10× linked reads | 1.03 Gb | Not reported / 715 | 137 kb | 3.3 Mb | 89.2% | Not reported |
| *Numida meleagris* (Galliformes) | [55] | Paired end + Mate pair | 1.04Gb | Not reported / 2,739 | 234 kb | 7.8 Mb | 90.7% | 19.5% |
| *Colinus virginianus* (Galliformes) | [56] | Paired end + HiC reads | 866 Mb | Not reported **/1,512 | Not reported ** | 66.8 Mb | 90.8% | Not reported |
| *Eremophila alpestris* (Passeriformes) | [57] | Paired end + mate pair | 1.04 Gb | Not reported ** / 2,708 | Not reported** | 10.6 Mb | 94.5% | Not reported |
| *Grus nigricollis* (Gruiformes) | [58] | Nanopore + paired end | 1.33 Gb | 1,837/ NA* | 17.9 Mb | NA* | 97.7% | 8.1% |
| *Melanerpes aurifrons* (Piciformes) | This study | Nanopore + paired end | 1.30 Gb/ | 441 / NA* | 16.0 Mb | NA* | 92.6% | 25.8% |

## Conclusions

We assembled the genome of the Golden-fronted Woodpecker (*Melanerpes aurifrons*), only the second published woodpecker genome. Our final assembly is 1.31 Gb in only 441 ungapped contigs with a contig N50 of 16 Mb. *k*-mer based estimates of genome size suggest that this assembly represents 95% of the actual genome, while a 92.6% recovery of bird-specific BUSCO genes suggests our assembly is relatively accurate. This genome should be foundational for studies of molecular evolution in woodpeckers and allies, and should also be useful for studies of woodpecker biodiversity, including phylogenetics, phylogeography, and establishing species limits in *Melanerpes*.

## Availability of Supporting Data and Materials

The genome assembly and raw reads were deposited to NCBI (project PRJNA598863).

## Acknowledgements

Promethion sequencing was carried out at the DNA Technologies and Expression Analysis Cores at the UC Davis Genome Center, supported by NIH Shared Instrumentation Grant 1S10OD010786-01. We thank OMRF for providing access to their computation cluster for data analysis. Scientific collecting was authorized by the Texas Parks and Wildlife Department and the United States Fish and Wildlife Service and supported by the Ornithology Department at the Sam Noble Museum. We thank them for their support of scientific collecting. We thank Joe Manthey, Texas Tech University, for discussions of woodpecker genomics.

## Author contributions

Conceptualization, benchwork, bioinformatics, writing original draft, review and editing: G.W., M.JM.; funding, voucher sample collecting: M.J.M.

---

**Supplementary Figure S1:**
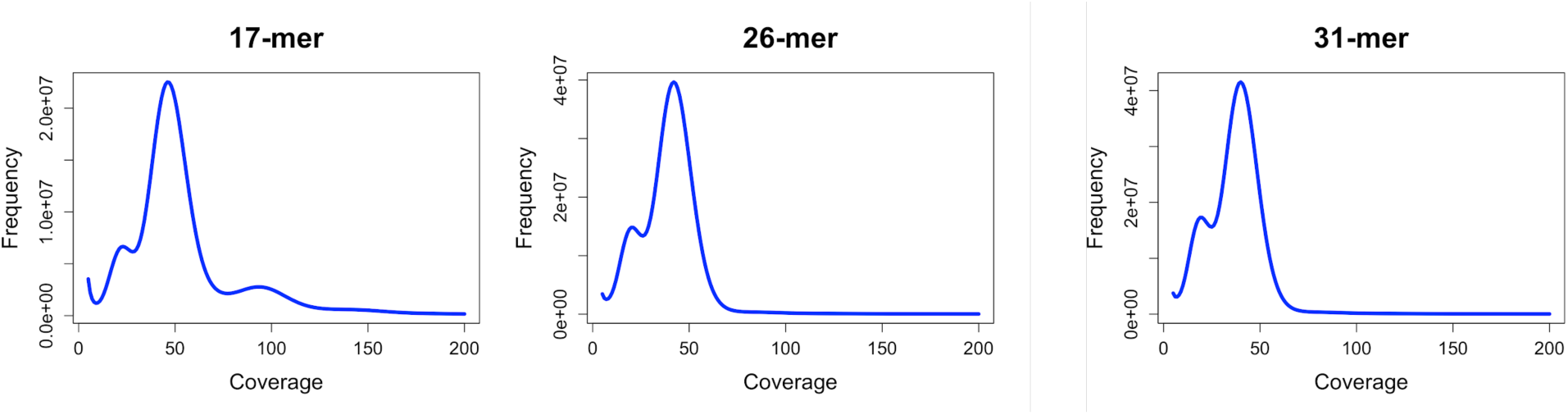
*k*-mer coverage distribution in our trimmed Illumina short reads for various *k*-mer values.

